# METTL3 promotes cell cycle progression via activation of transcriptional elongation

**DOI:** 10.1101/2025.07.12.664541

**Authors:** Azime Akçaöz-Alasar, Carlos A. Ramírez Parrado, Daniela Villalobos, Yanhua Wang, Prajwal C. Boddu, Alexander H. Nguyen, Hengyi Li, Marcelo Perez-Pepe, Pénélope Darnat, Yansheng Liu, Manoj M. Pillai, Lilian Kabeche, Claudio R. Alarcón

## Abstract

Transcription elongation through the phosphorylation of RNA Pol II by CDK9 (P-TEFb) is not only required for mRNA transcription to produce proteins, but is a fundamental mechanism required for proper cell cycle progression. N6-methyladenosine (m^6^A) methyltransferase METTL3 methylates mRNAs co-transcriptionally, modulating their downstream processing as well as the non-coding RNA 7SK post-transcriptionally. We found that CDK1 phosphorylates METTL3 at Ser43 at the onset of mitosis. This METTL3 phosphorylation promotes transcriptional elongation via the m^6^A/7SK/P-TEFb axis, allowing RNA Pol II clearance and proper cell cycle progression. Disruption of this pathway results in defects in chromosome segregation. Our findings establish a novel mechanistic link between CDK1 regulation of transcriptional elongation through activation of METTL3 and RNA methylation.

## Introduction

Transcription elongation is a fundamental regulatory step in cell cycle progression, ensuring the timely and coordinated expression of genes required for each phase of the cycle. At the onset of mitosis, global transcription initiation is largely inhibited, resulting in a “runoff” of actively elongating RNA polymerase II (RNAPII) and a subsequent cessation of transcriptional activity. This process is tightly coupled with chromatin restructuring and the removal of RNAPII from chromatin during the G2/M transition, which is essential for chromosome condensation and segregation (Bhowmick et al., 2023; Chan et al., 2012; Zhang et al., 2021).

The positive transcription elongation factor b (P-TEFb), a complex composed of CDK9 and cyclin T, is central to this regulatory network. P-TEFb phosphorylates the C-terminal domain (CTD) of RNAPII, as well as negative elongation factors such as NELF and DSIF, thereby releasing RNAPII from promoter-proximal pausing and enabling productive elongation (Fujinaga et al., 2023; Peterlin and Price, 2006). Early in mitosis, activation of P-TEFb is required for converting paused RNAPII into its elongating form, which is then able to clear gene bodies and dissociate from chromatin, enforcing the transcriptional shutdown necessary for accurate chromosome segregation and mitotic entry (Hirose and Ohkuma, 2007; Lu et al., 2016). Liang and colleagues showed that P-TEFb activation triggers the clearance of RNAPII complexes already engaged with chromatin, facilitating the transcriptional shutdown at the beginning of mitosis (Lee et al., 2025; Liang et al., 2015).

Disruption of these processes can lead to improper retention of RNAPII, aberrant gene expression during mitosis, and defects in cell division. Thus, the impact and regulation of transcription elongation—particularly via P-TEFb—are fundamental to maintaining cell cycle fidelity and genomic integrity (Fujinaga *et al*., 2023; Peterlin and Price, 2006).

N6-methyladenosine (m^6^A) is the most abundant internal modification in eukaryotic mRNA, and METTL3/METTL14, the enzymatic complex responsible for this RNA methylation, are implicated in the regulation of RNA stability (Wang et al., 2014a), miRNA processing (Alarcón et al., 2015b; Knuckles et al., 2017), RNA splicing (Alarcón et al., 2015a; Xiao et al., 2016), and translation (Lin et al., 2016; Meyer et al., 2015). These molecular processes regulate cellular functions such as meiosis (Clancy et al., 2002; Hongay and Orr-Weaver, 2011), cell proliferation (Fei et al., 2020; Lin *et al*., 2016; Wang et al., 2014b), and embryonic stem cell differentiation (Batista et al., 2014; Wang *et al*., 2014b), and pathophysiological states such as cancer (Cui et al., 2017; Xiang et al., 2017). Recently, m^6^A has been recognized as a pivotal regulator of gene expression (Akhtar et al., 2021; Perez-Pepe et al., 2023). We discovered that growth factors, such as EGF, activate MAPK signaling to induce the phosphorylation of METTL3 at position S43 by ERK. The phosphorylation of METTL3 induced its release from the sequestering factor HEXIM1, resulting in the METTL3-mediated methylation of 7SK. The m^6^A methylation of 7SK results in the displacement and release of the HEXIM/P-TEFb complex by HNRNP proteins. The free transcription elongation factor complex P-TEFb, in turn, phosphorylates the CTD of RNAPII, allowing transcription elongation to occur (Perez-Pepe *et al*., 2023).

Here we show that METTL3 is required for proper cell cycle progression. We find that CDK1 phosphorylates METTL3 at Ser43, releasing P-TEFb from its inactive complex with 7SK, and allowing transcription elongation to proceed. Depleting METTL3 from cells or preventing CDK-mediated phosphorylation results in accumulation of cells in G2/M and lagging and misaligned chromosomes. This phenotype is rescued by depletion of HEXIM1to release P-TEFb in an m^6^A-independent manner. These findings position METTL3/m^6^A as regulators of cell cycle progression through control of transcriptional elongation.

## Results

### METTL3 is required for cell proliferation and cell cycle progression

We and others have observed that depletion of METTL3 affects cell proliferation (Figures 1A and 1B). Disruption in several processes can result in defects in cell proliferation, including alterations in growth factor signaling, DNA replication/repair, protein synthesis, metabolism, apoptosis, etc. Among the main ones are defects in cell cycle progression. Thus, we decided to test if METTL3 depletion could result in cell cycle defects. We depleted METTL3 using two independent shRNAs (Figure 1A) and found that a decrease in the levels of METTL3 protein not only affect cell proliferation but results in accumulation of cells in G2/M as measured by DAPI flow cytometry in HeLa cells (Figures 1C and 1D) and the breast cancer cell line MDA-MB-231 (Supp. Figure 1). These results suggest that METTL3 is required for proper cell cycle progression.

**Figure 1.**
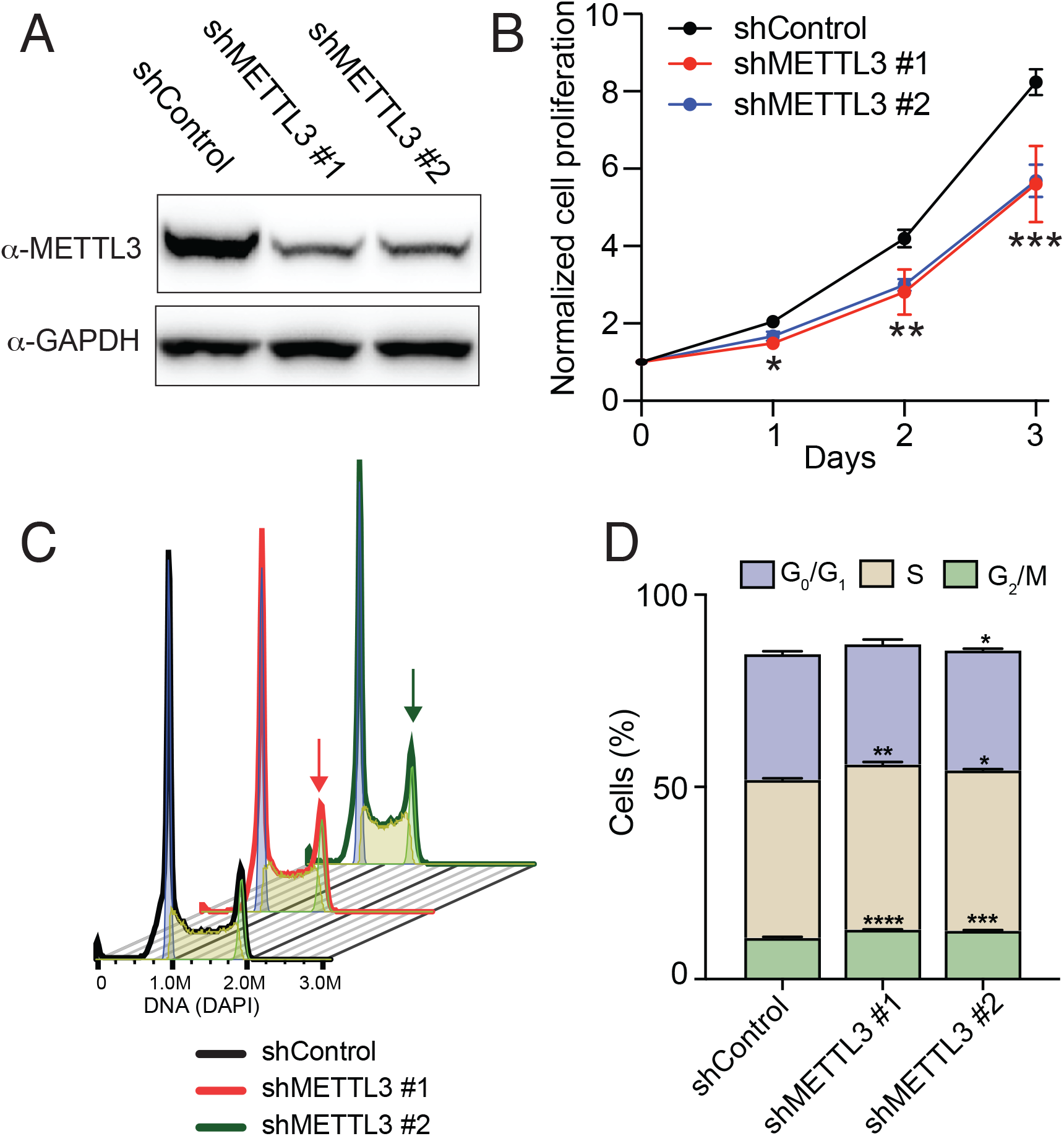
METTL3 depletion suppresses cell proliferation and promotes G2/M arrest. (**A**) METTL3 depletion using two independent shRNAs was compared to a shControl cell line and assessed by Western blot analyses using the indicated antibodies. (**B**) Proliferation rate of METTL3-depleted compared to control cells. (**C-D**) Flow cytometry analyses of cell cycle distribution in METTL3-depleted cells compared to control cells. A representative analysis (**C**) and the analysis of three independent experiments (**D**) are shown. Data are represented as mean ± SD (n = 3). Two-tailed Student’s t-test was performed to determine the statistical significance. *p ≤ 0.05, **p ≤ 0.01, ***p ≤ 0.001, ****p ≤ 0.0001.

### CDK1 phosphorylates METTL3 in early mitosis

We have recently found that growth factor signaling, including epidermal growth factor (EGF), results in the phosphorylation of METTL3 at position Ser43 (pS43) (Perez-Pepe *et al*., 2023). This phosphorylation results in the activation of METTL3 and, subsequently, the methylation of the non-coding RNA 7SK, which in turn is recognized by the RNA binding proteins HNRNPs to displace the inhibitory protein HEXIM1 and the positive transcription elongation factor P-TEFb (CDK9/CycT), finalizing with the phosphorylation of the RNAPII to promote transcription elongation. This was the first report connecting growth factor signaling to transcription through METTL3/m^6^A/7SK and the first one establishing a role for METTL3 as a transcriptional regulator in addition to its post-transcriptional roles (Perez-Pepe *et al*., 2023).

Since the cell cycle is a highly regulated process that involves sequential and orchestrated activation of multiple proteins, and we discovered that METTL3 can be regulated via phosphorylation, we hypothesize that the role of METTL3 in cell cycle progression could also be regulated in the same way. Thus, we analyzed our previous work, where we treated human U2OS cells either with nocodazole for 16 hours or left them untreated (control) (Rosenberger et al., 2017). Nocodazole is a microtubule-disrupting agent that primarily causes cell cycle arrest at the G2/M transition by blocking cells in mitosis. This is due to the inability to correctly assemble the mitotic spindle, thereby activating the spindle assembly checkpoint and halting cell cycle progression (Zieve et al., 1980). We acquired ten biological replicates, processed in parallel, each for nocodazole-treated and control samples, in Data-Independent Acquisition (DIA) mode, a bottom-up mass spectrometry approach that provides complete information on precursor and fragment ions (Chapman et al., 2014). We found that cells treated with nocodazole accumulated high levels of phosphorylated METTL3, an impressive more than 12-fold induction in pS43 compared to control cells (Figures 2A and 2B). To determine if the effect was due to the prolonged nocodazole treatment (16 hours), which could lead to DNA damage, we tested whether decreasing the time of exposure to nocodazole would result in similar pS43 phosphorylation. As shown in Figure 2C, short nocodazole treatment resulted in similar pS43 phosphorylation in HeLa cells, suggesting that METTL3 phosphorylation is likely not due to the deleterious effects induced by long nocodazole exposure.

**Figure 2.**
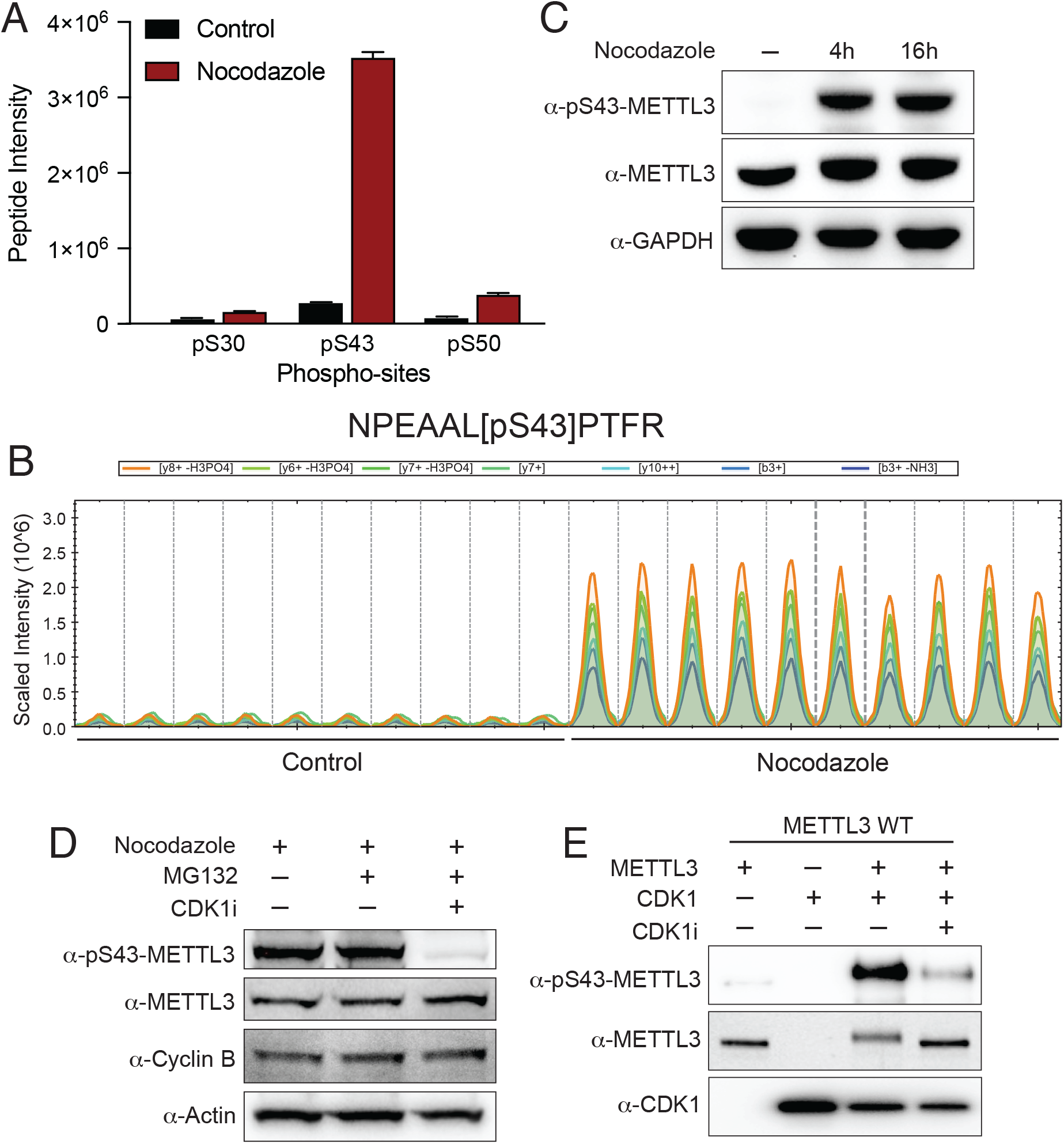
CDK1 phosphorylates METTL3 in early mitosis. (**A**) Bar graph representing the intensity of the indicated phospho-peptide as measured by mass spectrometry. Ten samples treated with nocodazole for 16 hours and ten control samples were analyzed. (**B**) Report the DIA-MS peaks (MS2 levels) of the samples described in A. (**C**) Western blot analyses of pS43-METTL3 in nocodazole-treated cells for indicated times. Total cell extracts were blotted, then subjected to the indicated antibodies. Blots are a representative of three biological replicates. (**D**) Effect of a CDK1 inhibitor RO3309 on the nocodazole-induced METTL3 phosphorylation. MG132 was used to stabilized Cyclin B. Total cell extracts were blotted, then subjected to the indicated antibodies. Blots are a representative of three biological replicates. (**E**) *In vitro* CDK1 kinase assay on purified METTL3 protein. Western blot was done using the indicated antibodies.

While we were searching for regulatory phosphorylation modulating METTL3 activity, we did not expect to find the same ERK-mediated phosphorylation that controls the methylation of 7SK (Perez-Pepe *et al*., 2023). Thus, we tested whether kinases controlling cell cycle progression could be responsible for the accumulation of pS43 in G2/M. The Ser43 is in a motif potentially recognized by kinases of the CMGC group (<u>C</u>DKs, <u>M</u>APKs, <u>G</u>SKs, and <u>C</u>LKs) (Suppl. Figure 2). Since CDKs are the kinases responsible for cell cycle control, we tested the possibility that CDK1, the master regulator of the CDKs, is the kinase that phosphorylates METTL3. Figure 2D shows that the CDK1 inhibitor RO3309, prevents the nocodazole-induced METTL3 phosphorylation. Moreover, an *in vitro* kinase assay using purified CDK1/Cyclin B on METTL3 protein shows that CDK1 can phosphorylate METTL3 at Ser43 *in vitro* (Figure 2E).

Taken together, these results show that METTL3 gets phosphorylated at Ser43 at G2/M as shown by nocodazole treatment, and that CDK1 is the likely kinase responsible for this phosphorylation.

### Phosphorylation of METTL3 at Ser43 is required for RNA Pol II clearance during early mitosis

To determine whether the pS43 phosphorylation in METTL3 has a functional role in cell cycle progression, we used CRISPR/Cas9 to mutate Ser43 to alanine (S43A) as we have done previously (Perez-Pepe *et al*., 2023). As shown in Figure 3A, two independent S43A clones were insensitive to EGF-stimulated pS43 phosphorylation without affecting METTL3 expression level, thus validating the specificity of the mutation. The S43A clones mimic the cell proliferation defects (Figure 3B) and the cell cycle accumulation in G2/M (Figures 3C and 3D) observed when METTL3 was depleted, demonstrating that the effect of METTL3 in proliferation and cell cycle progression depends on the CDK-mediated phosphorylation at Ser43.

**Figure 3.**
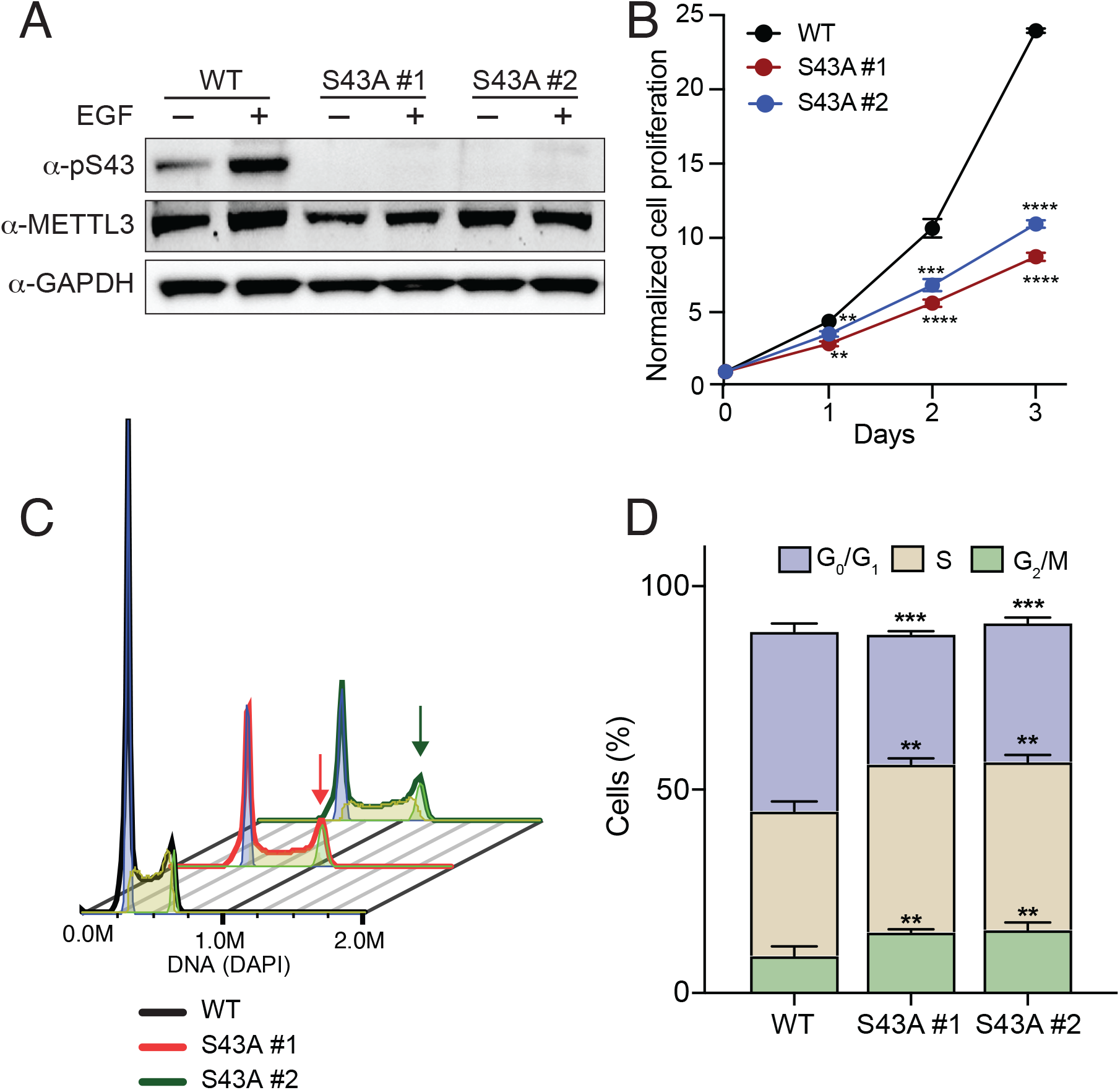
METTL3 phosphorylation is required for proper cell proliferation and cell cycle progression. (**A**) Western blot analyses of pS43-METTL3 in EGF-stimulated WT cells and two clones carrying a S43A METTL3 mutation. (**B**) Cell proliferation rate of cells shown in A. Each data point represents the average of three biological replicates. (**C**) Representative example of flow cytometry analyses of DAPI-stained cells, analyzed by FlowJo software. (**D**) Quantification of cell cycle phases distribution in three biological replicates of WT and S43A mutant cells. Data were analyzed by the two-tailed t-test to determine the significance among groups at each time point. **p ≤ 0.01, ***p ≤ 0.001, ****p ≤ 0.0001.

Since we just established the functional role of pS43 METTL3 in cell cycle progression, and we previously showed the role of pS43 in transcription elongation, the logical question is to ask whether pS43 METTL3 is involved in cell cycle progression through the regulation of transcription elongation. It has been previously shown that P-TEFb activation triggers the clearance of RNAPII complexes already engaged with chromatin, facilitating the transcriptional shutdown at the beginning of mitosis (Liang *et al*., 2015). Thus, we decided to test the impact of the METTL3 S43A mutation on RNAPII clearance during the cell cycle using “Native elongation transcript sequencing using mammalian cells” (mNET-seq), which allows the examination of genome-wide nascent transcript profiles based on the phosphorylation status of RNAPII and coupled RNA processing (Nojima et al., 2016). To perform mNET-seq, we isolated S43A METTL3 and wild-type cells, either synchronized in G2/M by nocodazole treatment or asynchronous. To observe the effect of CDK9 (P-TEFb) on transcription elongation and clearance during mitosis, we additionally treated the cells with the CDK9 inhibitor Flavopiridol. Once the cells are collected, the chromatin fraction is isolated, and an antibody against phosphorylated RNAPII at position pSer2 is used for immunoprecipitation. The extracted RNA associated with DNA-bound RNAPII was extracted and prepared for RNA-seq analysis. As Liang and colleagues previously showed, RNAPII is bound to the promoter regions in asynchronous cells, but it is cleared in cells treated with nocodazole, i.e., accumulated in G2/M (Figures 4A and 4B). However, when cells are treated with nocodazole and Flavopiridol, thus inhibiting CDK9, then an increase in RNAPII bound to the promoter region sis observed. Figure 4C shows a heat map of the normalized read distribution of the samples, centered on the transcription start site (TSS). As we had hypothesized, mutation of S43A in METTL3 resulted in a defect of RNAPII to be cleared from nocodazole-treated cells, validating the role of pS43-METTL3 in releasing and activating P-TEFb to promote transcriptional elongation and clearance (Figure 4). These results demonstrate that the phosphorylation of METTL3 at Ser43 is functional and required for proper transcription elongation during cell cycle progression.

**Figure 4.**
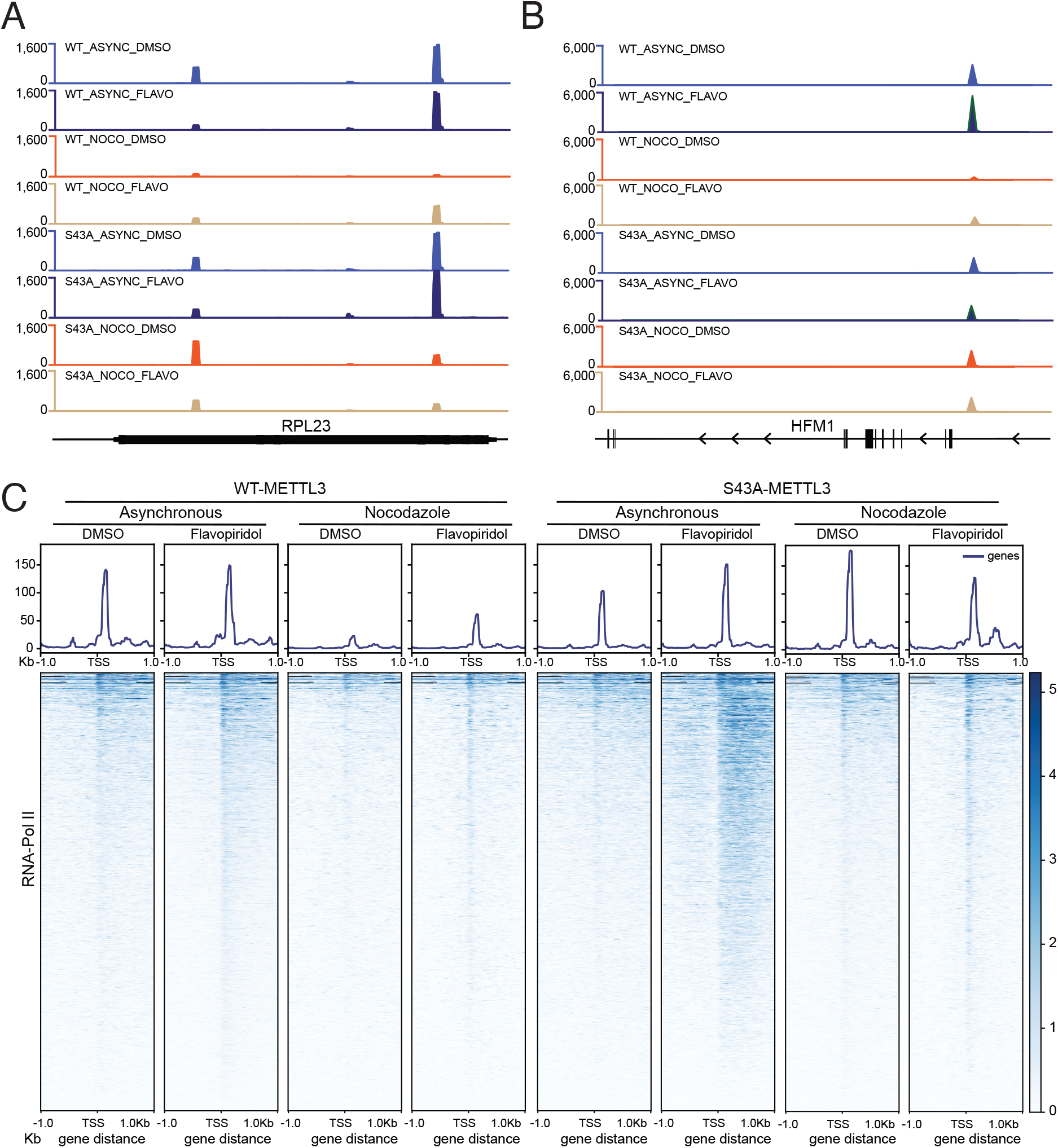
Elimination of METTL3 phosphorylation inhibits the release of promoter-proximal RNA Pol II in early mitosis. mNET-seq of pSer2-RNAPII was used to determine changes to transcription elongation. (**A-B**) Representative genome coverage plots of WT and S43A cells, asynchronous or synchronized with Nocodozaole, with or without CDK inhibition by Flavopiridol. (**C**) Heat map of normalized read distribution of same mNET-seq samples in A and B, centered on transcription start site (TSS) and 1 Kb upstream or downstream.

### HEXIM1 depletion restores proliferation and cell cycle progression induced by lack of METTL3 phosphorylation

Figure 5A depicts a mechanistic model of the role of pS43 METTL3 in transcription elongation. Phosphorylation of METTL3 is required for the methylation of 7SK and exchange of HEXIM1 for HNRNP proteins, resulting in the release of P-TEFb (Perez-Pepe *et al*., 2023). When S43 is mutated to alanine, METTL3 is not activated and can no longer methylate 7SK, thus keeping P-TEFb bound to 7SK through interaction with HEXIM1. If this model is correct, we could release P-TEFb by depleting HEXIM1, thus allowing transcription elongation and cell cycle progression in a METTL3/m^6^A-independent manner. Therefore, we knockdown HEXIM1 with two independent shRNAs in the context of S43A CRISPR/Cas9 mutation of METTL3, as shown in Figure 5B. HEXIM1 knockdown partially rescues the proliferation defect observed in the S43A-METTL3 mutant cells (Figure 5C). As anticipated, the depletion of HEXIM1 rescues the cell progression defect and returns the percentage of cells in G2/M cells to wild-type levels (Figures 5D and 5E).

**Figure 5.**
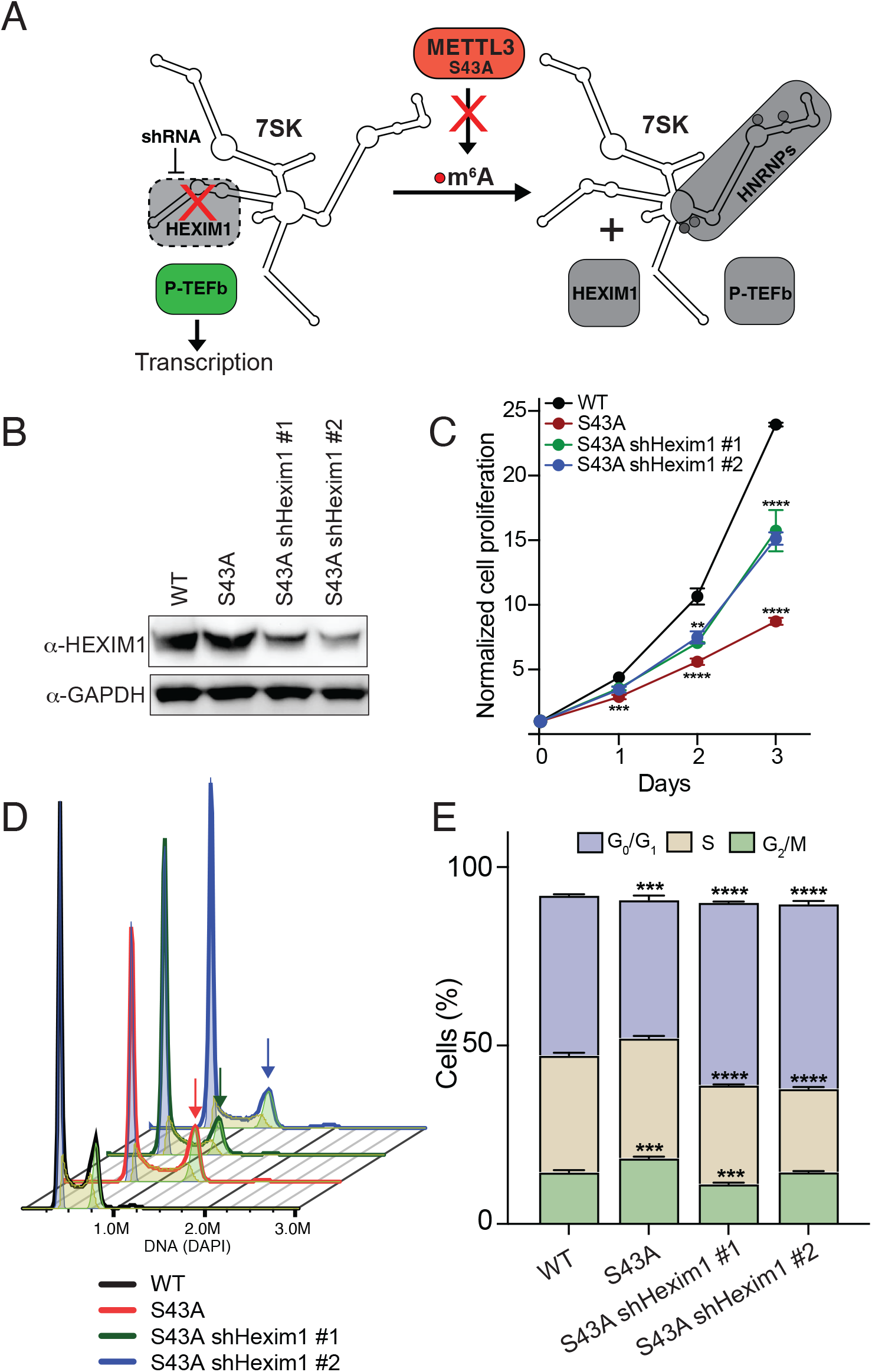
HEXIM1 prevents cell METTL3-dependent cell cycle progression. (**A**) Model depicting the effect of HEXIM1 elimination on transcription in the context of a S43A METTL3 mutant. Inactive members of the pathway are colored in gray. S43A METTL3 mutation results in the inhibition of 7SK methylation and failed to release P-TEFb. This effect could be rescue by depleting cells from HEXIM1 thus releasing P-TEFb independently of METTL3 methylation of 7SK. (**B**) Knockdown of HEXIM1 with two independent shRNAs. GAPDH was used as a loading control. (**C**) Cell proliferation rate of WT, S43A, and S43A shHEXIM1 cells. Data are presented as mean value ± standard deviation. (**D-E**) Cell cycle progression of WT, S43A, and S43A shHEXIM1 cells. A representative histogram of the indicated cells stained with DAPI (**D**) and the analysis of three independent experiments (**E**) are shown. Error bars represent mean ± SD of three biological replicates. Statistical analyses were carried out by a two-tailed t-test (**: p ≤ 0.01, ***: p ≤ 0.001, ****: p ≤ 0.0001).

These results support the model by which the axis CDK/METTL3/7SK/P-TEFb regulates transcription elongation required for cell cycle progression.

### Disruption of METTL3/m^6^A results in aberrant chromosome segregation

Numerous studies have established a strong link between cell cycle defects and the segregation of chromosomes, including lagging or misaligned chromosomes during mitosis. These errors can lead to improper chromosome segregation, genomic instability, and aneuploidy, which are hallmarks of many cancers and other diseases. (Potapova and Gorbsky, 2017; Thompson and Compton, 2011). Previously, it was found that Flavopiridol treatment increases the incidence of chromosome segregation defects, including lagging and misaligned chromosomes as well as anaphase bridges (Liang *et al*., 2015). Therefore, we analyzed the effect of METTL3 depletion on chromosome segregation. Cells were seeded in coverslips and 24 hours later fixed with paraformaldehyde, permeabilized, incubated with the primary antibodies, and stained with DAPI. Images for chromosome missegregation defects and chromosome analysis are collected and analyzed using both Nikon’s NIS-elements and Image J software. Figures 6A and 6B show representative images of misalignments and lagging chromosomes, respectively. Depletion of METTL3 significantly increased the percentage of both misalignments (Figure 6C) and lagging chromosomes (Figure 6D), indicating that the defects observed in cell progression upon METTL3 depletion have detrimental consequences on chromosomal segregation. We finally ask whether those effects could also be dependent on the pS43 phosphorylation of METTL3. Indeed, when we used S43A mutant METTL3 cells, those chromosome segregation defects were recapitulated (Figures 6E and 6F). Importantly, depletion of HEXIM1 in the context of the S43A mutation rescues the detrimental effects caused by this mutation (Figures 6E and 6F).

**Figure 6.**
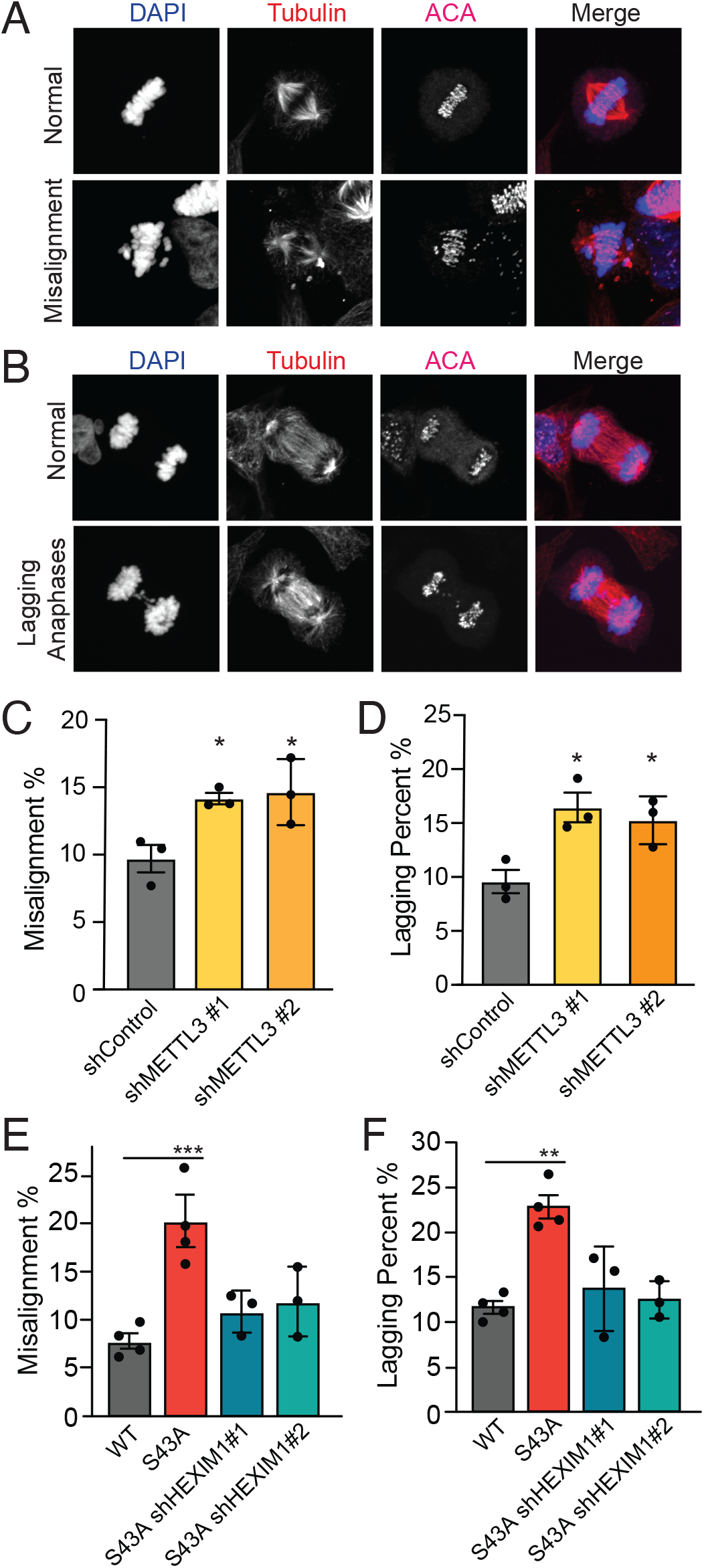
METTL3 dysregulation results in lagging and misaligned chromosomes. (A) Representative z-projection images of HeLa cells showing aligned or misaligned chromosomes. (B) Representative z-projection image of HeLa cells showing lagging chromosome defects. Bar graphs representing the quantification of misalignment rates (C) or lagging anaphase rates (D) when METTL3 is depleted with two independent shRNAs compared to a control shRNA. Bar graphs representing the quantification of misalignment rates (E) or lagging anaphase rates (F) when METTL3 carried a S43A mutation and the effect of HEXIM1 depletion using two independent shRNAs. Statistical analyses were carried out by a two-tailed t-test (*: p ≤ 0.05, **: p ≤ 0.01, ***: p ≤ 0.001).

Taken together, these results indicate that the CDK1-mediated pS43-METTL3-dependent transcription elongation and clearance of RNAPII are necessary for proper cell division, and alterations in this pathway may result in detrimental consequences to chromosomal segregation.

## Discussion

In this report, we uncover a fundamental aspect of METTL3/m^6^A biology, which is the regulation of cell cycle progression. Taking together the results presented here supports a model where CDK1 activates METTL3 through phosphorylation of Ser43 to induce the methylation of 7SK and release of P-TEFb to promote transcription elongation, RNAPII clearance, and cell cycle progression (Figure 7).

**Figure 7.**
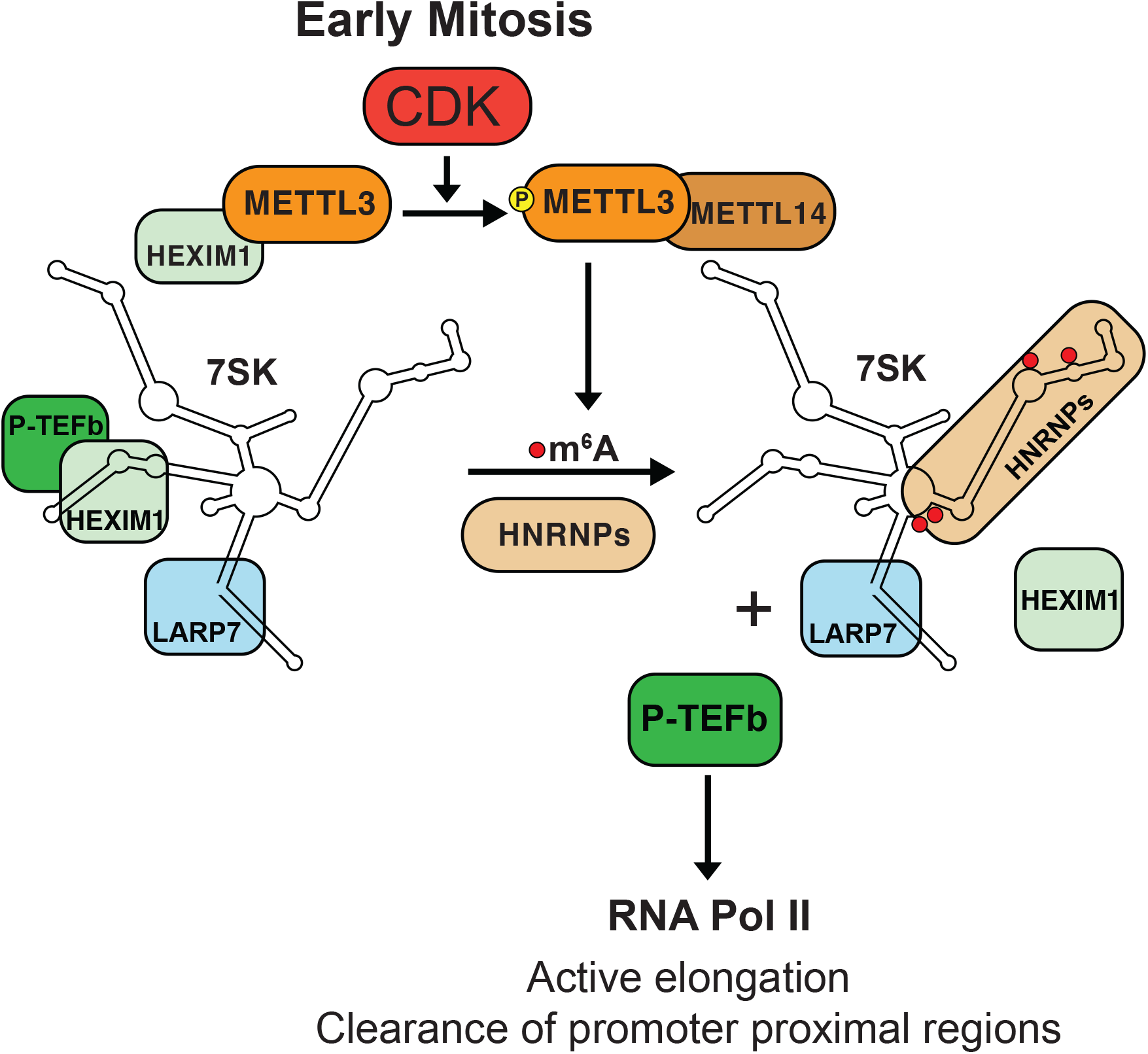
Model. The work presented here is summarized in this model. At the beginning of mitosis CDKs phosphorylate METTL3 at position S43. This phosphorylation activates METTL3 allowing the methylation of 7SK and release of P-TEFb complex. Free P-TEFb is then available to stimulate the release of paused Pol II to clear the promoter proximal region through transcription elongation allowing cell cycle progression.

This model is consistent with our previous discovery of METTL3 acting as a transcriptional regulator in addition to its more established post-transcriptional role. Both processes are fundamentally different; on one hand, post-transcriptional mRNA modification occurs co-transcriptionally in association with RNA Pol II (Knuckles *et al*., 2017), while the methylation of 7SK occurs in a mature, RNA Pol III non-coding RNA target (Perez-Pepe *et al*., 2023). It is important to note that while the mechanism of METTL3 activation via pS43 phosphorylation is the same under growth factor stimulation and cell cycle progression, the kinases responsible for this phosphorylation are different. Growth factor stimulation activates the MAPK signaling pathway, which results in METTL3 phosphorylation by ERK, while during the cell cycle, CDK1 is the kinase responsible for it. Both MAPKs and CDKs belong to the CMGC group, which is a well-conserved family of protein kinases distinguished by a common kinase core and a distinctive CMGC-insert segment, which contributes to their ability to recognize specific substrates. Members of this family include cyclin-dependent kinases (CDKs), mitogen-activated protein kinases (MAPKs), glycogen synthase kinases (GSKs), and CDC-like kinases (CLKs). These enzymes are integral to the regulation of essential cellular functions such as controlling the cell cycle, promoting cell proliferation, guiding differentiation, and apoptosis (Chowdhury et al., 2023). It is an open question whether additional members of the CMGC group could phosphorylate METTL3 under different conditions to use transcriptional regulation as the last step of the signaling cascade.

Xi and colleagues recently identified in cancer cells a link between m^6^A, glucose metabolism, ROS levels, and CDK2 activity, leading to cell cycle arrest or progression (Xi et al., 2025). It would be interesting to investigate is this mechanism applies to normal cells under metabolic stress or other conditions affecting glucose or reactive oxygen species.

Our findings identify METTL3/m^6^A as a modular component of the cell cycle, which is key for cell proliferation and proper chromosome segregation. Future studies are needed to define the complete array of biological processes modulated by the modification m^6^A.

## Methods

### Cell Culture

HeLa cells were maintained in DMEM (Gibco) supplemented with 10% fetal bovine serum (Gibco) and 1% penicillin-streptomycin (Gibco) in a humidified atmosphere of 5% CO_2_ at 37 °C. For EGF stimulation, 70% confluent cells were serum-starved for 16h, then incubated with 100 ng/ml recombinant human EGF protein (R&D Systems 236-EG-200) for 30 min. To prolong mitosis, the cells were treated with 10 µg/ml nocodazole for 4h and 16 h, then collected through mitotic shake-off and proceeded with protein extraction.

Mutant METTL3 S43A cell lines were previously generated by the CRISPR-Cas9 system (Perez-Pepe et al. 2023). shRNA-mediated knockdown cell lines were generated as described previously (Perez-Pepe et al. 2023). shRNAs inserted pLKO 2.0 plasmids were co-transfected with pCMV_VSVG and pCMV_dR.8 plasmids to human embryonic kidney 293T cells to produce lentivirus. Two shRNAs targeting METTL3 (GCAAGTATGTTCACTATGAAA and GCCAAGGAACAATCCATTGTT); two shRNAs targeting HEXIM1 (GCGGCATTGGAAACCGTACTA and GCGGCATTGGAAACCGTACTA), and a control shRNA (SHC002) were used.

### Western Blot

Cells were lysed using ice-cold RIPA buffer (Thermo Fisher Scientific) with protease inhibitors (Sigma-Aldrich) and phosphatase inhibitors (Roche). Cell lysates were passed through QIAshredder (QIAGEN). 25 µg total protein was resolved by electrophoresis on 4%–12% NuPAGE bis-tris gels and then transferred to nitrocellulose membranes (Bio-Rad). The membranes were blocked with 5% milk in TBS-T. Primary antibodies, rabbit HEXIM1 (12604, CST); mouse α-METTL3 (H00056339-B01P, Abnova); cyclin-B; CDK1; rabbit α-GAPDH horseradish peroxidase (HRP) conjugate (8884, CST), incubations were carried out in 5% milk or 5% BSA in TBS-T for either 1 h. at room temperature or overnight at 4°C. Polyclonal antibody against pS43 METTL3 was generated by Thermo Fisher Scientific. The membranes were incubated with secondary antibodies, goat α-rabbit HRP-linked (7074, CST); donkey α-mouse IgG H&L (IRDye 680RD) preabsorbed (ab216779, Abcam) in 5% milk in TBS-T (chemiluminescence) or 3% BSA (fluorescence), respectively. Subsequently, the membranes were developed using the ECL Plus reagent (Amersham), and chemiluminescent signals were captured on a Bio-Rad Chemidoc.

### Cell proliferation assay

Cell proliferation rate was measured using the XTT cell proliferation kit (Roche Applied Science) according to the manufacturer’s instructions. Briefly, 1 x10^4^ cells were seeded into 24-well plates. After 5h, 0.3 mg/ml XTT was added to the cells and incubated at 37°C for 2 h to obtain at 0 h measurement. The absorbance at 480 nm and 660 nm was read in a plate reader (BioTek Synergy H1 Plate Reader). The protocol was repeated at 24 h, 48 h, and 72 h. The values were normalized to the 0 h measurement.

### Cell cycle

70-80% confluent cells were harvested by trypsinization, fixed in 70% ethanol, and stored at -20°C °C overnight. The cells were pelleted and washed with PBS. Finally, cells were pelleted again and resuspended in 0.6 mg/ml DAPI in 0.1% (v/v) Triton X-100 (prepared in PBS). They were incubated for 30 minutes at RT, protected from light. The cells were analyzed with CyTek Aurora (CyTek Bioseciences) flow cytometry, and the distributions of cell cycle phases were calculated using FlowJo software.

### Nascent RNA-seq

mNET-seq was carried out as previously described (Nojima *et al*., 2016). In brief, the chromatin fraction was isolated from 2.5×10^7^ HeLa cells for each sample. Then the chromatin was digested in 100 μL of MNase (40 units/ μL) reaction mixture for 2 min at 37°C in a thermomixer (1,400 rpm). 1.25ul EGTA (1M) was added and incubated on ice for 10min to inactivate MNase, and the soluble digested chromatin was collected after 16,000 rpm centrifugation for 10 min at 4°C. The supernatant was diluted with 900 μL of NET-2 buffer (50 mM Tris-HCl pH 7.4, 150 mM NaCl and 0.05% NP-40, RNase inhibitor) and Pol II antibody-conjugated beads were added. The Immunoprecipitation was performed at 4°C for 1.5 hr. And 5 μg of RNAPII CTD phospho-Ser2 mAb (ACTIVE&MOTIF 61083) was used per sample. After immunoprecipitation, the beads were washed with 1 mL of NET-2 buffer 8 times and washed with 300 μL of 1xPNKT (1xPNK buffer and 0.05% Triton X-100) buffer once in the cold room. Then resuspended the beads in 50 μL PNK reaction mix (1xPNKT, 1.5 mM ATP and 1U/ul T4 PNK 3′phosphatase minus (NEB), inhibitor) and incubated in Thermomixer (1,400 rpm) at 37°C for 6 min. After the reaction beads were washed with 1 mL of NET-2 buffer once and RNA was extracted with 200ul TRIzol reagent and 0.5 ng of DNAse-treated Drosophila RNA spike-ins were added to each sample. RNA was suspended in 10ul urea Dye (7M Urea, 1xTBE, 0.1% BPB and 0.1% XC) and resolved on 8% TBU gel (Invitrogen) at 30w 30min. The 30-100 nt RNAs were cut between BPB and XC dye markers. The small RNAs were eluted from crushed gel using 400ul RNA elution buffer (1 M NaOAc and 1 mM EDTA) by rotating at 25°C for 2 hr. Eluted RNA was purified with a Spin-X column and the flow-through RNA was ethanol precipitated. mNET-seq libraries were prepared with Qiagen smRNA library prep kit. The libraries were sequenced on an Illumina NovaSeq S4 (paired-end 2 × 150) at an average depth of 100 million reads per sample.

### Immunofluorescence

Cells were seeded and grown for 1 day prior until 70-80% confluency in a 12-well culture-treated plate with coverslips. Then, the cell media was removed, and a fixing solution of 3.5% paraformaldehyde was added for 15 min. Subsequently, cells were washed with phosphate buffered saline (PBS) solution, permeabilized with Triton X-100 (PBS+1% + Triton X-100) for 10 min, washed 5 min in PBS solution and finally blocked with BSA (PBS +BSA 1%) for 20 min to 1 hour. Each well was incubated overnight at 4 °C with 500 μL of primary antibody diluted in BSA (PBS+BSA 1% + 1:1000 primary antibody). The coverslips were then washed with PBS for 5 minutes and incubated for 1 hour at room temperature with secondary antibody solutions (PBS+BSA 1% + 1:1000 secondary antibody) containing DAPI staining (1mg/mL). A final wash with PBS is performed for 5 minutes, and then coverslips are extracted from the plate to be mounted on microscopy slides using ProLong Gold Antifade Mountant (Invitrogen). Mounted slides are covered from light and left to dry overnight at room temperature. Images for chromosome missegregation defects and chromosome analysis are collected on a Nikon ECLIPSE Ti W2 spinning disk for confocal microscopy with a Hamamatsu Fusion sCMOS camera. Image were analyzed using both Nikon’s NIS-elements and Image J (Fiji) software.

### *In vitro* kinase assays and recombinant proteins

Human CDK1/Cyclin B HIS-Tag protein was purchased from Invitrogen (PV3292), and GST fusion-METTL3 was expressed and purified from bacteria cells. DMSO was used as a loading control in each reaction at the equivalent volume used during drug-inhibited conditions. CDK1/Cyclin B during inhibited conditions was performed by adding CDK1 inhibitor RO3309 at a concentration of 5 uM diluted in DMSO. The in vitro kinase assays were conducted in a kinase assay buffer (20 mM 3-morpholinopropane-1-sulphonic acid (MOPS), pH 7.2, 25 mM MgCl_2_, 500 μM ATP, 1 mM DTT) at 30 °C while shaking gently. Each reaction contained 250 nM of each protein according to the reaction condition. Quenching of the reactions was achieved by adding 5 mM EDTA, and the samples were processed for subsequent immunoblotting.

