## Supplementary figures and images for "METTL3 promotes cell cycle progression via activation of transcriptional elongation"

### Suppl. Figures

#### Supplemental Figure 1

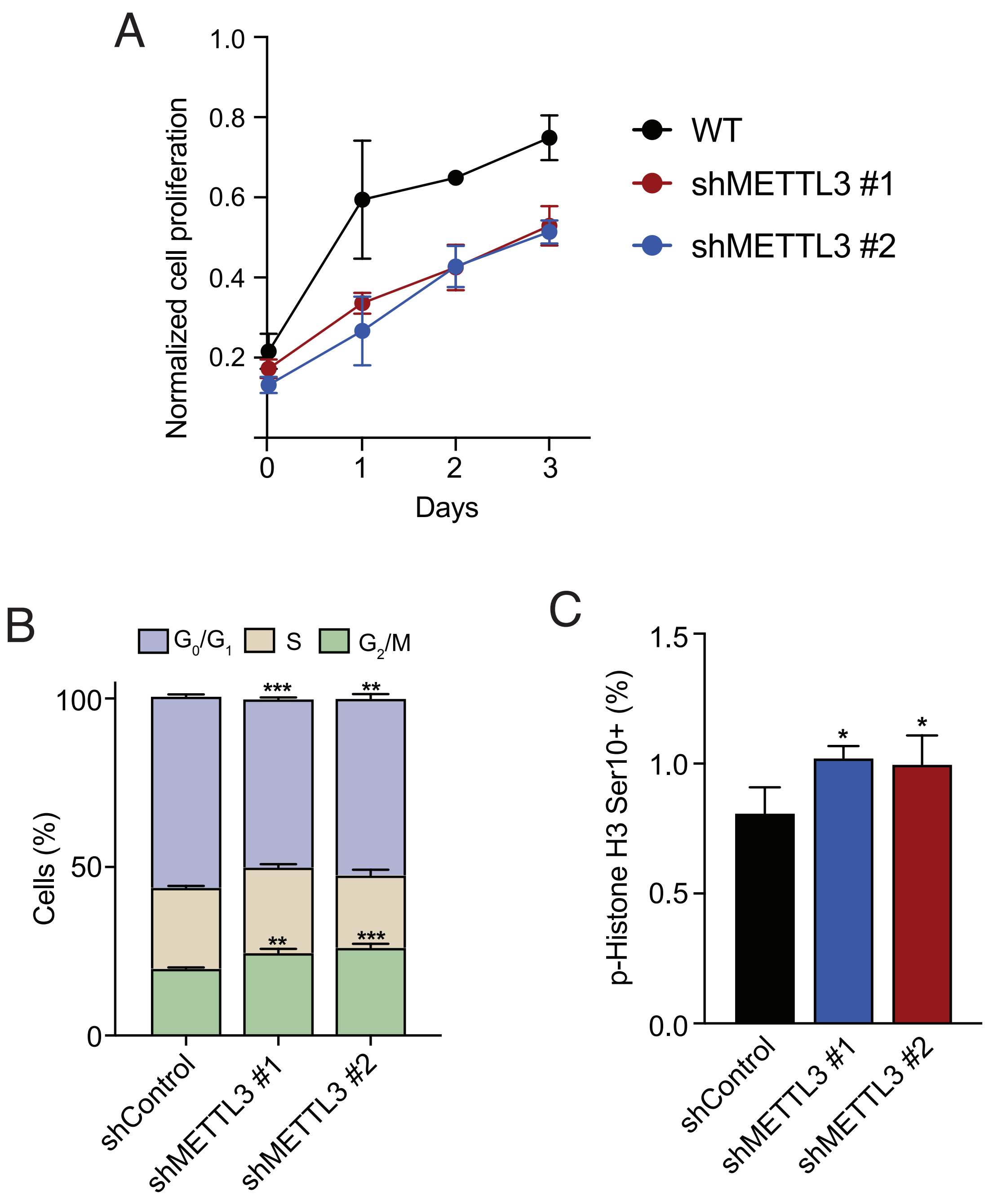

#### Supplemental Figure 2

pS43-Predicted Kinases

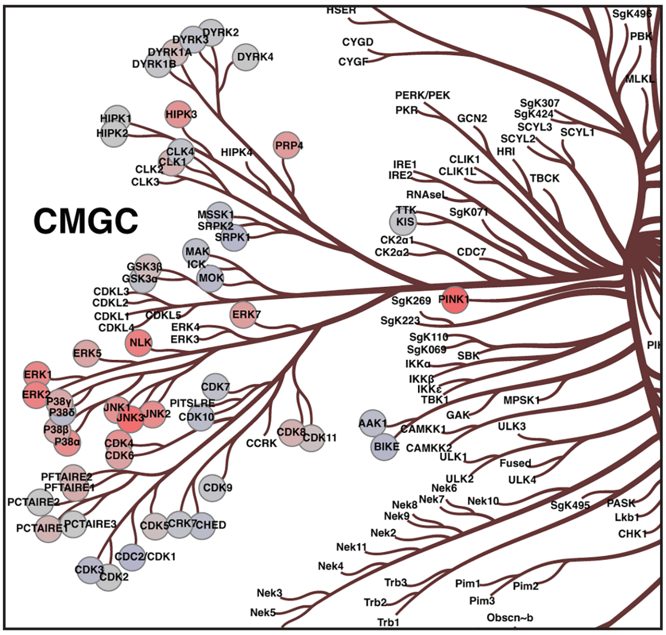
